# Triglyceride metabolism controls inflammation and *APOE4*-associated disease states in microglia

**DOI:** 10.1101/2024.04.11.589145

**Authors:** Roxan A. Stephenson, Kory R. Johnson, Linling Cheng, Linda G. Yang, Jessica T. Root, Jaanam Gopalakrishnan, Han-Yu Shih, Priyanka S. Narayan

## Abstract

Microglia modulate their cell state in response to various stimuli. Changes to cellular lipids often accompany shifts in microglial cell state, but the functional significance of these metabolic changes remains poorly understood. In human induced pluripotent stem cell-derived microglia, we observed that both extrinsic activation (by lipopolysaccharide treatment) and intrinsic triggers (the Alzheimer’s disease-associated *APOE4* genotype) result in accumulation of triglyceride-rich lipid droplets. We demonstrate that lipid droplet accumulation is not simply concomitant with changes in cell state but rather necessary for microglial activation. We discovered that both triglyceride biosynthesis and catabolism are needed for the transcription and secretion of proinflammatory cytokines and chemokines in response to extrinsic stimuli. Additionally, we reveal that triglyceride biosynthesis and catabolism are necessary for the activation-associated phagocytosis of multiple substrates including the disease-associated amyloid-beta peptide. In microglia harboring the Alzheimer’s disease risk *APOE4* genotype, triglyceride-rich lipid droplets accumulate even in the absence of any external stimuli. Inhibiting triglyceride biosynthesis in *APOE4* microglia not only modifies the transcription of immune response genes but also attenuates disease-associated transcriptional states. This work establishes that triglyceride metabolism is necessary for microglia to respond to extrinsic activation. In *APOE4* microglia, this metabolic process modulates both immune signaling and a disease-associated transcriptional state. Importantly, our work identifies metabolic pathways that can be used to tune microglial immunometabolism in *APOE4-*associated disease.

## Introduction

Neuroinflammation and disruptions to lipid metabolism have emerged as central to the pathogenesis and progression of many neurodegenerative diseases, including Alzheimer’s disease (AD). Microglia, the resident immune cells of the CNS, are primary mediators of neuroinflammation; they adopt a variety of proinflammatory states in response to activating stimuli including pathogens, injury, protein aggregates, and cellular degeneration. Recent studies have characterized the multitude of microglial transcriptional states in mouse models, stem-cell derived human microglia, and human post-mortem brain tissue^1–7^. Taken together, these studies clarify that microglia are plastic and can adopt a myriad of states in response to their environment. These states can impact signaling of other cell types and contribute to disease^8–10^. Microglia highly express many genes harboring single nucleotide polymorphisms that are enriched in AD patients relative to controls, further implicating microglia in the pathogenesis and progression of AD^11^.

Metabolic disruptions are emerging as central to the biology of several neurodegenerative diseases including AD. Genome-wide association studies implicate genes in lipid metabolism including the apolipoprotein genes *APOE* and *CLU,* the lipid phosphatase, *INPP5D*, lipid signaling gene, *PLCG2,* and lipid transporters, ABCA7 and ABCA1, among others^12^. Intriguingly, glial lipid accumulation was one of the original pathological hallmarks noted by Dr. Alois Alzheimer in 1907^13^. In addition, recent work in AD mouse models and AD patient samples display lipid droplet accumulation within various glial cell types including microglia^14,15^. The *APOE4* allele is associated with a 3-fold (heterozygous) or a 12-fold (homozygous) increase in AD incidence^16^. As a lipid transport protein, *APOE4* has been associated with the accumulation of cholesterol and triglycerides in several model systems^17,18,27,19–26^. Our previous work identified that *APOE4* increases triglyceride storage in lipid droplets in human induced pluripotent stem cell (iPSC)-derived astrocytes and microglia^22^. This has now been observed in multiple other systems including neuron-microglial co-cultures^23,24^, post-mortem brain samples ^24,28^, and mouse models^17,27,29^. Recent reports even suggest that *APOE* functions as a lipid-droplet binding protein^30^.

A growing body of research has revealed a complex interplay between the metabolic state of innate immune cells and their response to environmental stimuli. In microglia, inflammatory stimuli have been shown to shift metabolism from reliance on oxidative phosphorylation to glycolysis^31^. In aged mice, lipid droplets accumulate in microglia, which also exhibit a proinflammatory state^32^. Challenging mouse microglia with aggregates of the AD-associated amyloid-beta peptide also elicits lipid droplet accumulation^15,19,24^. In both human and mouse microglia, *APOE4*-mediated changes to microglial lipid metabolism, including lipid droplet accumulation, are correlated with a proinflammatory state^19,20,23,24^. Altering the metabolic state of microglia can have cell non-autonomous consequences on neuronal communication^23,24^. Lipid droplet accumulation in microglia due to exposure to amyloid-beta aggregates or the *APOE4* genotype can cause downstream effects on neuronal calcium signaling and neuronal tau phosphorylation^15,24^. Taken together, these studies have strongly correlated microglial lipid droplets with inflammatory functions. However, we still do not understand whether lipid droplets are an essential part of the inflammatory response or an epiphenomenon. We also do not understand the similarities and differences between lipid droplet accumulation initiated by extrinsic versus intrinsic stimuli.

In this work, we define the connections between microglial triglyceride metabolism and response to inflammatory stimulation, both extrinsic and intrinsic. We used human iPSC-derived microglia which can be either activated using compounds such as lipopolysaccharide or exhibit an immune active state when they harbor disease genotypes like *APOE4*. When stimulated by external ligands or when harboring disease genotypes, these microglia display accumulation of triglyceride-rich lipid droplets. We manipulated triglyceride biosynthesis and catabolism in activated and resting microglia with chemical inhibitors to discover that both synthesis and catabolism of triglyceride-rich lipid droplets are necessary for the microglial response to extrinsic stimulation. Moreover, we show that triglyceride biosynthesis can modulate the immune and disease-associated responses of microglia harboring the AD-associated *APOE4* genotype. Taken together, our work demonstrates that modulating triglyceride metabolism with chemical inhibitors can tune microglial responses to both extrinsic and intrinsic immune stimuli.

## Results

### LPS-activated microglia display neutral lipid accumulation

Microglia accumulate triglycerides upon extrinsic immune activation. To understand the mechanisms connecting microglial lipid metabolism and inflammation, we used human iPSC-derived microglia, which allow us to contextualize our findings in the setting of human health and disease. This is especially important since glial lipid metabolic pathways have been shown to differ between mouse and human^20^. To accomplish this objective, we utilized a well-characterized iPSC cell line with an *APOE3* homozygous genotype^21^ and normal stable karyotype (Figure S1A). We used established protocols^33^ to derive human iPSCs into microglia (Figure S1C) which were CD11b positive by FACS (Figure S1D), and expressed canonical microglial markers IBA1 and CX3XR1 (Figures 1A, S1E). These cells did not express canonical markers of other neural lineage cell types (Figure S1E).

**Figure 1.**
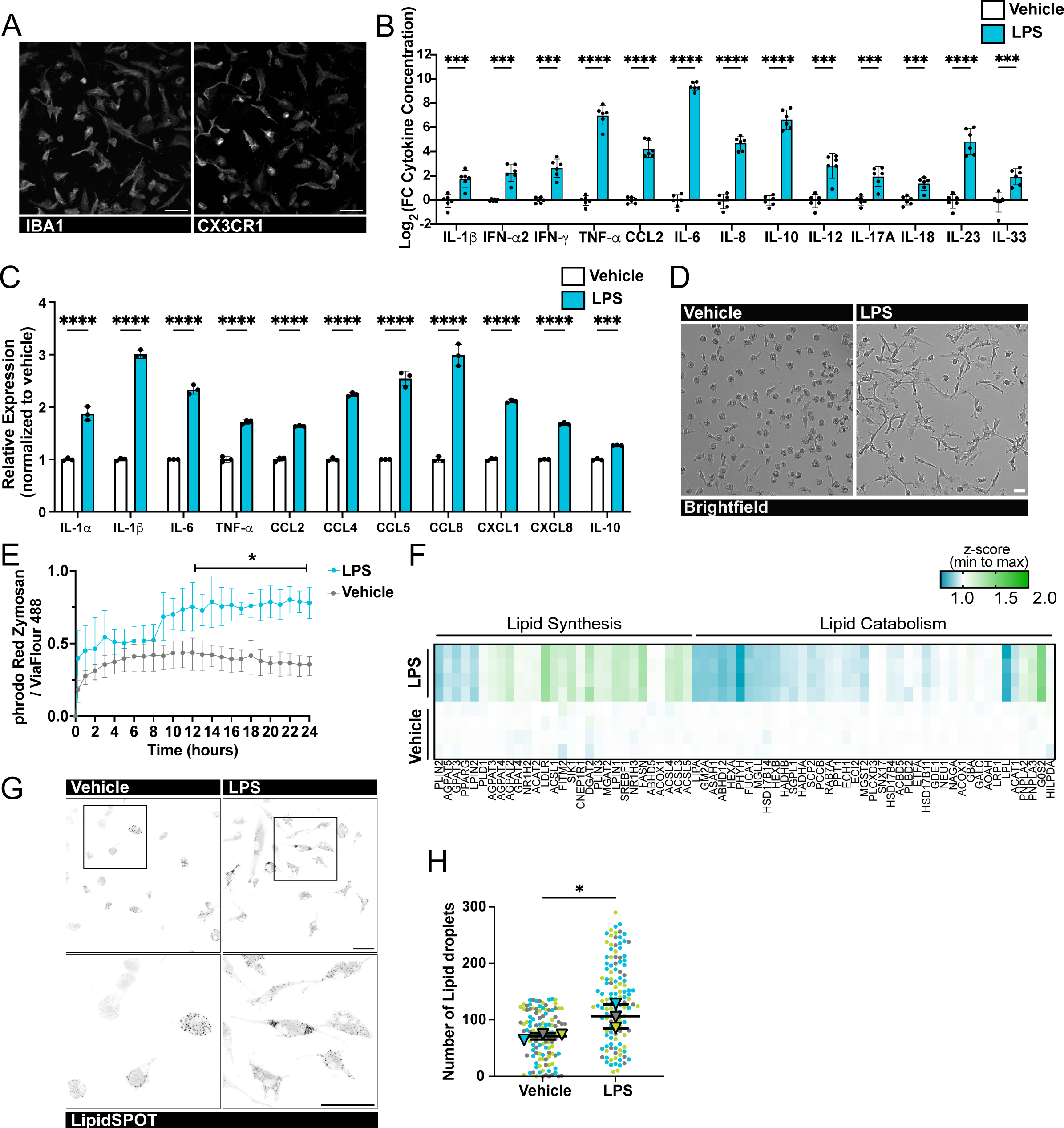
Neutral lipid homeostasis is affected during microglia activation. **A.** Fluorescence images of *APOE3/APOE3* human iPSC-derived microglia stained with antibodies against microglia markers, IBA1 (left panel) and CX3CR1 (right panel). Scale bar: 50 μm. **B.** Quantification of the fold change of cytokines secreted by vehicle-treated (white bars) and lipopolysaccharide (LPS)-treated (blue bars) *APOE3/APOE3* microglia. n = 6 wells for 3 distinct biological cultures. *** P ≤ 0.001; **** P ≤ 0.0001 by unpaired t-tests. **C.** Quantification of the expression levels of cytokines (from normalized read counts) relative to average of vehicle-treated controls (vehicle, white bars; LPS, blue bars; n = 3). **** P ≤ 0.0001 by two-way ANOVA, with post-hoc Šídák. **D.** Representative brightfield images of vehicle-treated (right) and LPS-treated (left) *APOE3/APOE3* microglia. Scale bar: 100 μm. **E.** A 24h time course of fluorescence intensity of pHrodo Red Zymosan normalized to the cytoplasmic stain, ViaFlour 488, quantified per cell over 24 hours (vehicle, gray; LPS, blue; n = 3 wells). * P ≤ 0.05 by unpaired t-tests. **F.** Heatmap of differentially expressed genes associated with lipid synthesis and lipid catabolism. **G.** Representative fluorescence images of *APOE3/APOE3* microglia stained with the neutral lipid dye, LipidSPOT. Scale bars: 50 μm. High magnification images (bottom panels) show regions of interest in top panels. **H.** Quantification of the lipid droplet number, with each dot representing a cell. Each color represents an independent microglia differentiation. Triangles depict the average number of lipid droplets in each biological replicate. * P ≤ 0.05 by unpaired t-test.

We then confirmed that these human iPSC-derived microglia were able to respond to immune stimuli by treating them with lipopolysaccharide (LPS). After 16h of LPS treatment, the microglia secreted increased levels of proinflammatory cytokines and chemokines into the cell culture media (Figure 1B) and increased their expression of several genes known to be involved in immune activation (Figure 1C). The iPSC-derived microglia exhibited a profound morphological change upon LPS-mediated activation (Figure 1D) as well as increased phagocytosis of fluorescent zymosan-functionalized bioparticles (Figure 1E).

When we investigated the broader effects of LPS stimulation, transcriptomic analysis showed that activated microglia upregulate genes associated with lipid synthesis and downregulate many genes associated with lipid catabolism (Figure 1F), suggesting increased accumulation of cellular lipids. These transcriptomic changes were associated with increased accumulation of neutral-lipid rich lipid droplets (Figure 1G,H). We ensured that these changes to lipid accumulation upon LPS activation occurred independently of the cell’s genetic background and derivation protocol by testing microglia derived from another iPSC line, WTC-11 (Figure S1F), using an orthogonal microglial differentiation protocol^34^ (Figure S1G). In this second line, we observed similar accumulation of lipids upon LPS activation (Figure S1H), demonstrating that LPS-mediated lipid accumulation is robust across genetic backgrounds and derivation protocols.

### Triglyceride biosynthesis is necessary for LPS-mediated activation of microglia

Our transcriptomic data revealed that LPS-mediated microglial activation induced changes in multiple fatty acid and triglyceride biosynthesis enzymes (Figure 1F). Excess fatty acids are often stored in triglyceride-rich lipid droplets, which we observed by microscopy (Figure 1G,H). Other studies have observed increased levels of microglial lipid droplets in the contexts of aging, disease, and inflammation ^15,23,24,27,32^, suggesting that lipid accumulation in microglia is a central biological phenomenon and operates in multiple contexts. To further understand the functional consequences of lipid accumulation we chose to interrogate whether lipid accumulation was necessary for microglial activation or whether it was an epiphenomenon.

In this study, we focused on the triglyceride metabolism pathway. We took advantage of well-characterized and highly specific inhibitors of the enzymes DGAT1 and DGAT2 (diacylglycerol acid transferases 1 and 2) which control the rate limiting step of triglyceride biosynthesis from diacylglycerol and fatty acids (Figure 2A). In our transcriptional data, *DGAT2* expression increased upon LPS-mediated activation (Figure 2B). Although *DGAT1* expression did not increase with LPS activation (Figure 2B), we chose to inhibit both enzymes simultaneously to ensure that functional redundancy between the enzymes did not confound our data ^35^. We performed a live/dead assay and confirmed that this compound treatment did not impact cell viability (Figure S2A,B). We established an experimental paradigm where we first inhibited triglyceride biosynthesis and then stimulated the microglia with LPS prior to downstream assays (Figure 2C). Upon treatment of our hiPSC-derived microglia with a combination of DGAT1 and DGAT2 inhibitors (DGAT inhibitor), we observed an expected reduction in triglyceride lipid accumulation even when stimulated with LPS (Figure 2D,E). This reinforces the fact that, in microglia, LPS treatment stimulates triglyceride biosynthesis via DGAT activity.

**Figure 2.**
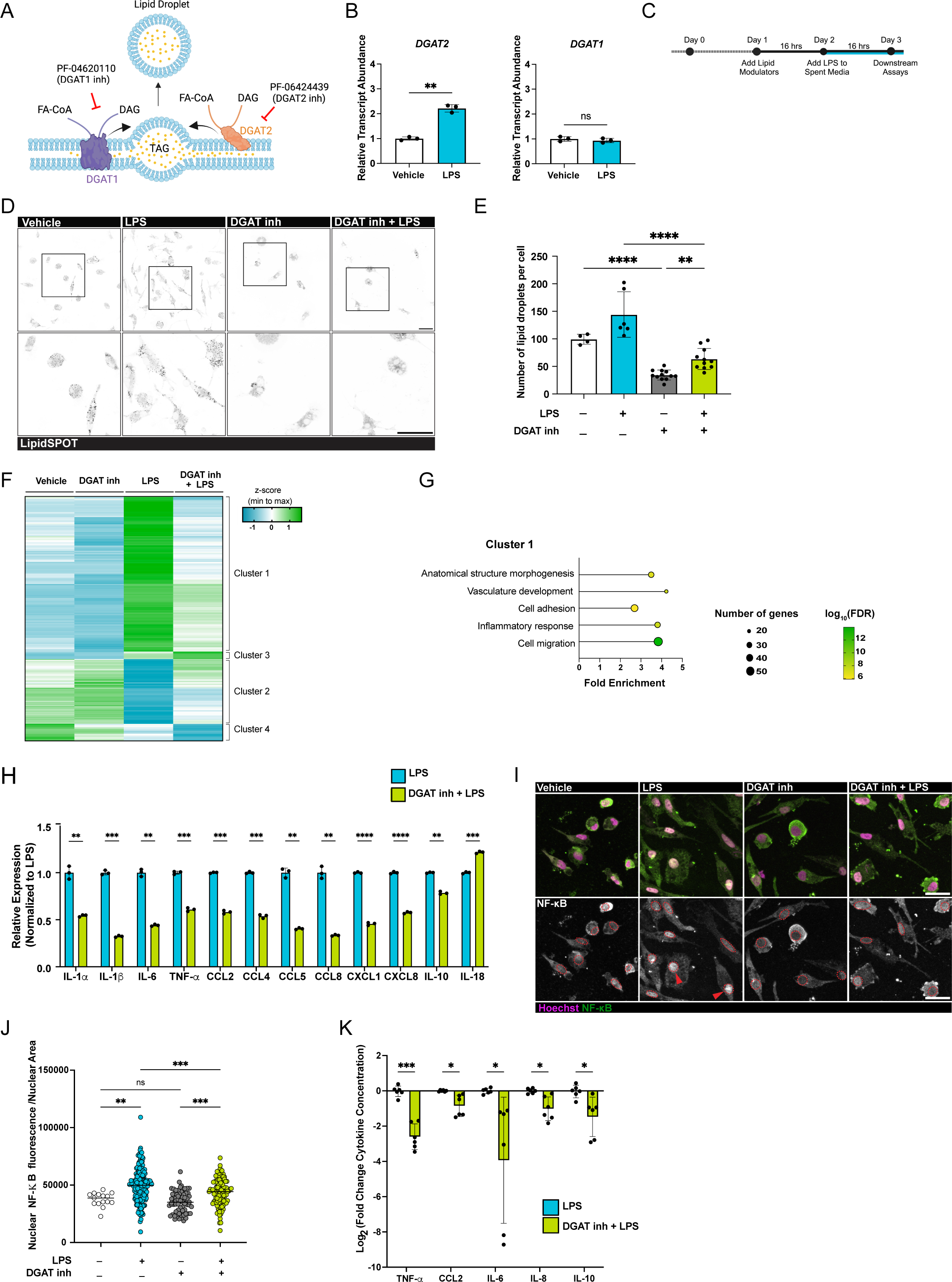
Triglyceride accumulation and LD-biogenesis are necessary for LPS-mediated activation.

Following DGAT inhibitor treatment, we stimulated the microglia with LPS and profiled their transcriptomes. DGAT inhibitor treatment in the context of LPS activation, drove a distinct transcriptional program from DGAT inhibitor treatment alone, which had little effect. We subdivided this transcriptional response into four clusters of genes which respond differently to LPS in the presence and absence of DGAT inhibitor treatment (Figure 2F). Most differentially regulated genes belonged to cluster 1, which contained genes that were upregulated upon LPS treatment but downregulated when treated with both DGAT inhibitor and LPS. When we analyzed the gene ontology terms associated with this downregulation, they encompassed not only the expected lipid metabolism genes (such as *ACSL1*, *DGAT2*, *FASN*, *CREB5*, *MBOAT7*, and *AGPAT4*) but also many involved in the inflammatory response (like *IL-6, IL-1b, TNF-a, CXCL5, CXCL8, PTGES*, and *PTGS2*) (Figure 2G, S2C). When we interrogated a panel of cytokine and chemokine genes that were increased in expression upon LPS treatment, we observed that DGAT inhibitor treatment inhibited this upregulation (Figure 2H). Notably, in the absence of LPS stimulation, microglia treated with DGAT1 and DGAT2 inhibitors showed no significant changes in the expression of inflammatory genes suggesting context-specific regulation (Figure S2D).

Since we observed the differential regulation of many inflammatory genes, we wanted to know whether NF-κB, one of the primary transcriptional regulators of inflammation, was impacted. NF-κB translocates from the cytosol to the nucleus upon inflammatory activation. LPS treatment resulted in a robust translocation of NF-κB of microglia to the nucleus whereas LPS treatment in the presence of DGAT inhibitor treatment reduced the number of microglia with nuclear-localized NF-κB (Figure 2I,J). This finding suggested that DGAT inhibition is important for processes upstream to inflammatory gene transcription like NF-κB translocation. We asked whether the large changes to gene expression upon LPS and DGAT inhibitor treatments were due to alterations in chromatin accessibility. When we performed ATAC sequencing (assay for transposase-accessible chromatin with sequencing), we observed that LPS treatment resulted in an expected increase in chromatin accessibility at several inflammatory genes (Figure S2E, Sy.x=16.11). Although DGAT inhibitor treatment decreased the expression of many genes upregulated upon LPS treatment, it did not impact chromatin accessibility within the body of these genes (Figure S2F, Sy.x=9.107). This result leads us to believe that DGAT inhibitor treatment acts downstream of chromatin remodeling.

When we inhibited triglyceride biosynthesis and then activated the microglia using LPS, we noticed decreased levels of secreted cytokines and chemokines compared to when microglia were treated with LPS alone (Figure 2K, S2G). In the absence of LPS activation, DGAT inhibitor treatment did not impact the cytokine secretion in microglia (Figure S2H). We had observed that LPS-mediated activation resulted in a change in microglial morphology (Figure 1D). DGAT inhibitor treatment prevented this change in morphology (Figure S2I,J). This phenotype was consistent in microglia derived from iPSCs of another healthy genetic background (Figure S1B, S2K,L). Taken together, these data reveal that inhibition of triglyceride biosynthesis directly impacts the ability of microglia to respond to inflammatory stimuli like LPS, establishing that triglyceride biosynthesis is necessary for inflammatory activation of microglia.

Upon LPS-mediated activation, microglia exhibit increased phagocytosis (Figure 1E). We asked whether inhibition of triglyceride biosynthesis would impact the downstream effects of microglial activation on phagocytosis. Upon DGAT inhibitor treatment followed by LPS stimulation, we no longer observed the LPS-associated increase in phagocytosis of zymosan bioparticles (Figure 3A). Decreased uptake upon DGAT inhibitor treatment (in the context of LPS stimulation) was also observed for other ligands as well—fluorescent dextrans which enter via clathrin-mediated pathways (Figure 3B,C) and amyloid-beta, the aggregating peptide associated with AD (Figure 3D,E). These effects on amyloid-beta phagocytosis held true in microglia derived from iPSCs of a different healthy genetic background (Figure S1B, S3A,B). We also noticed that DGAT inhibitor treatment in the presence of LPS resulted in lower expression of *FPR1* and *FPR2*, two genes involved in microglial phagocytosis, when compared to LPS alone (Figure S3C). This finding supports a hypothesis that the effects of DGAT inhibitor treatment on activation-associated phagocytosis may occur at the transcriptional level. Taken together, our investigation of phagocytosis reveals that inhibition of triglyceride biosynthesis can not only impact the inflammatory response to LPS but also can modulate cellular uptake of disease-relevant substrates.

**Figure 3.**
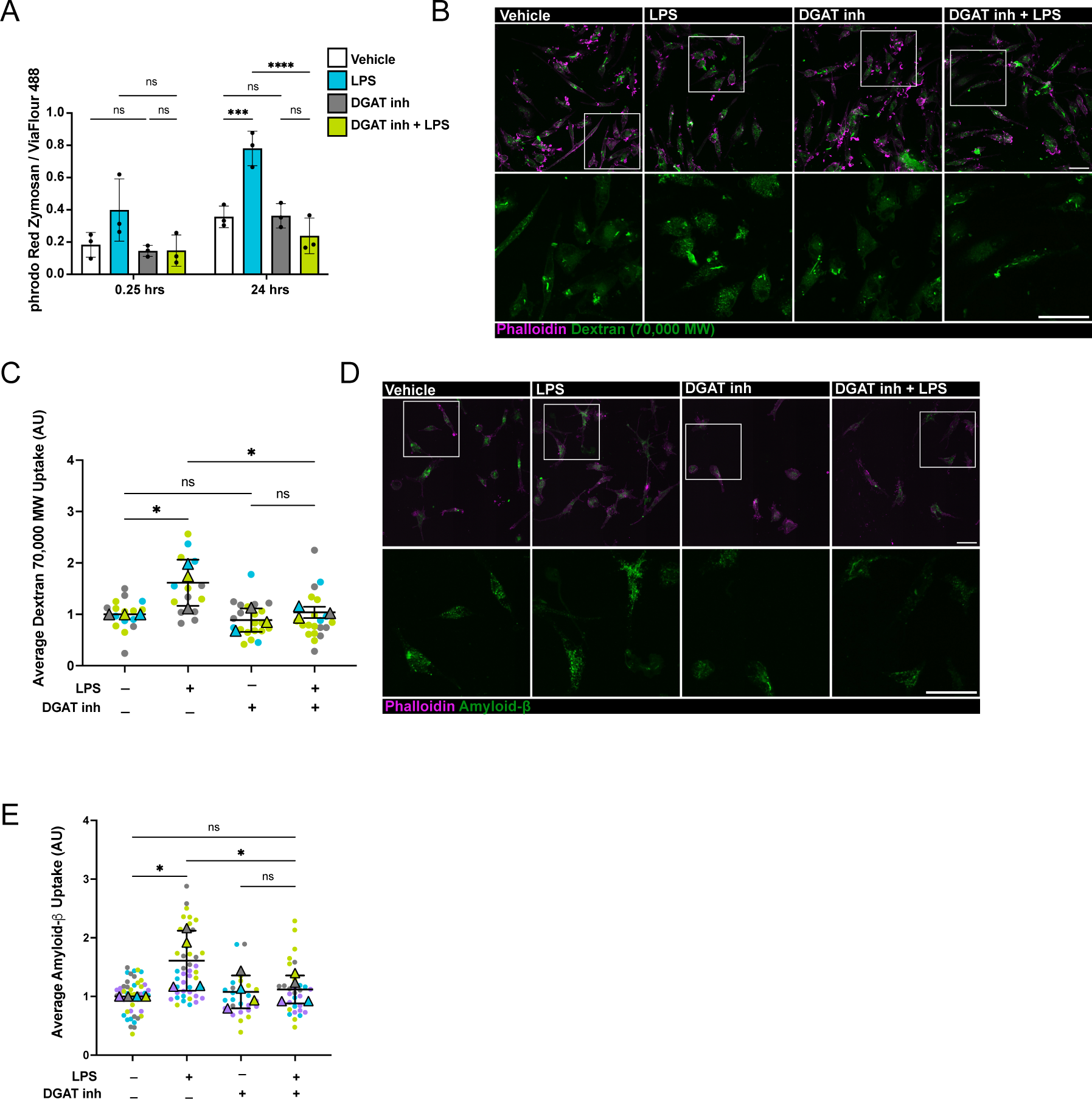
Modulating triglyceride accumulation and LD-biogenesis alters uptake in microglia. **A.** Quantification of the fluorescence intensity of pHrodo Red Zymosan uptake normalized to ViaFlour 488 cytoplasmic stain per cell at time = 0.25 hours and time = 24 hours (Vehicle, white; LPS-treated, blue; DGAT1 and DGAT2 inhibition, gray; DGAT1 and DGAT2 inhibition + LPS; green; n = 3 wells). ns P > 0.05; *** P ≤ 0.001; **** P ≤ 0.0001 by one-way ANOVA, with post-hoc Šídák test to correct for multiple comparisons. **B.** Representative fluorescence images of dextran (green) uptake assay in resting and activated *APOE3/APOE3* microglia in the presence or absence of DGAT1 and DGAT2 inhibitors. Cells were stained with Phalloidin (magenta). Scale bars: 50 μm. Bottom panels show higher magnification of white box in top panels. **C.** Quantification of fluorescent dextran uptake assay (B) relative to vehicle-treated controls. Each dot represents an average of at least 20 cells per image. Each color represents an independent microglia differentiation. Triangles depict the average dextran uptake in each biological replicate. ns P > 0.05; * P ≤ 0.05 by one-way ANOVA, with post-hoc Šídák test to correct for multiple comparisons. **D.** Representative fluorescence images of amyloid-beta (green) uptake Assay in homeostatic and activated *APOE3/APOE3* microglia in the presence or absence of DGAT1 and DGAT2 inhibitors. Cells were stained with Phalloidin (magenta). Scale bars: 50 μm. Bottom panels show higher magnification of white box in top panels. **E.** Quantification of amyloid-beta uptake (D) relative to vehicle-treated controls. Each dot represents an average of at least 20 cells per image. Each color represents an independent microglia differentiation. Triangles depict the average amyloid-beta uptake in each biological replicate. ns P > 0.05; * P ≤ 0.05 by one-way ANOVA, with post-hoc Šídák test to correct for multiple comparisons.

We specifically chose to target DGAT1 and DGAT2 due to their rate limiting role in triglyceride biosynthesis. However, we also explored whether inhibition of enzymes further upstream in the triglyceride biosynthesis pathway have similar effects. ACSL1 (acyl-coenzyme A synthase long chain family member 1) catalyzes the conjugation of a coenzyme A group to fatty acid acyl chains to enable the synthesis of many lipid types and has recently been identified as upregulated in the context of AD^24^. We observed an upregulation of ACSL1 when treating our healthy microglia with LPS (Figure S3D). We observed that inhibition of ACSL1 reduced the number of cellular lipid droplets in a similar manner to DGAT1 and DGAT2 inhibition (Figure S3E,F). Additionally, ACSL1 inhibition also had a similar effect to DGAT1 and DGAT2 inhibition on zymosan bioparticle phagocytosis in the context of LPS activation (Figure S3G), suggesting that targeting flux through the triglyceride biosynthesis pathway further upstream similarly impacts the microglial response to LPS.

### Triglyceride catabolism is necessary for LPS-mediated activation of microglia

We established that triglyceride synthesis is necessary for a complete microglial response to LPS. We then asked whether the presence of triglyceride-rich lipid droplets alone was sufficient for activation of microglia by LPS. To investigate this further, we used chemical inhibitors of ATGL (adipose triglyceride lipase) and DDHD2 (DDHD domain containing 2), two enzymes that catalyze the first step of triglyceride catabolism from lipid droplets (Figure 4A). Upon LPS activation, we found that *ATGL* but not *DDHD2* gene expression was increased suggesting that ATGL may play a role in microglial activation (Figure 4B, S4A). Inhibition of either of these enzymes (ATGL or DDHD2) resulted in increased lipid droplets in our human iPSC-derived microglia (Figure 4C,D, S4B,C). We then measured the effects of these inhibitors on secreted cytokines levels in the presence or absence of LPS mediated activation. Neither ATGL inhibitor nor DDHD2 inhibitor treatment altered secreted cytokine levels in unactivated cells (Figure S4D,E) revealing that although triglyceride-rich lipid droplets are necessary for activation, they are not sufficient to elicit a strong activation response from microglia. However, when ATGL inhibitor or DDHD2 inhibitor treatment was followed by LPS stimulation, we noticed reduced secretion of multiple cytokines compared to LPS treatment alone (Figure 4E, S4F,G). The effect of ATGL inhibitor treatment on cytokine secretion far exceeded that of DDHD2 inhibitor treatment, revealing that triglyceride catabolism via ATGL is necessary for microglia to fully respond to extrinsic activation.

**Figure 4.**
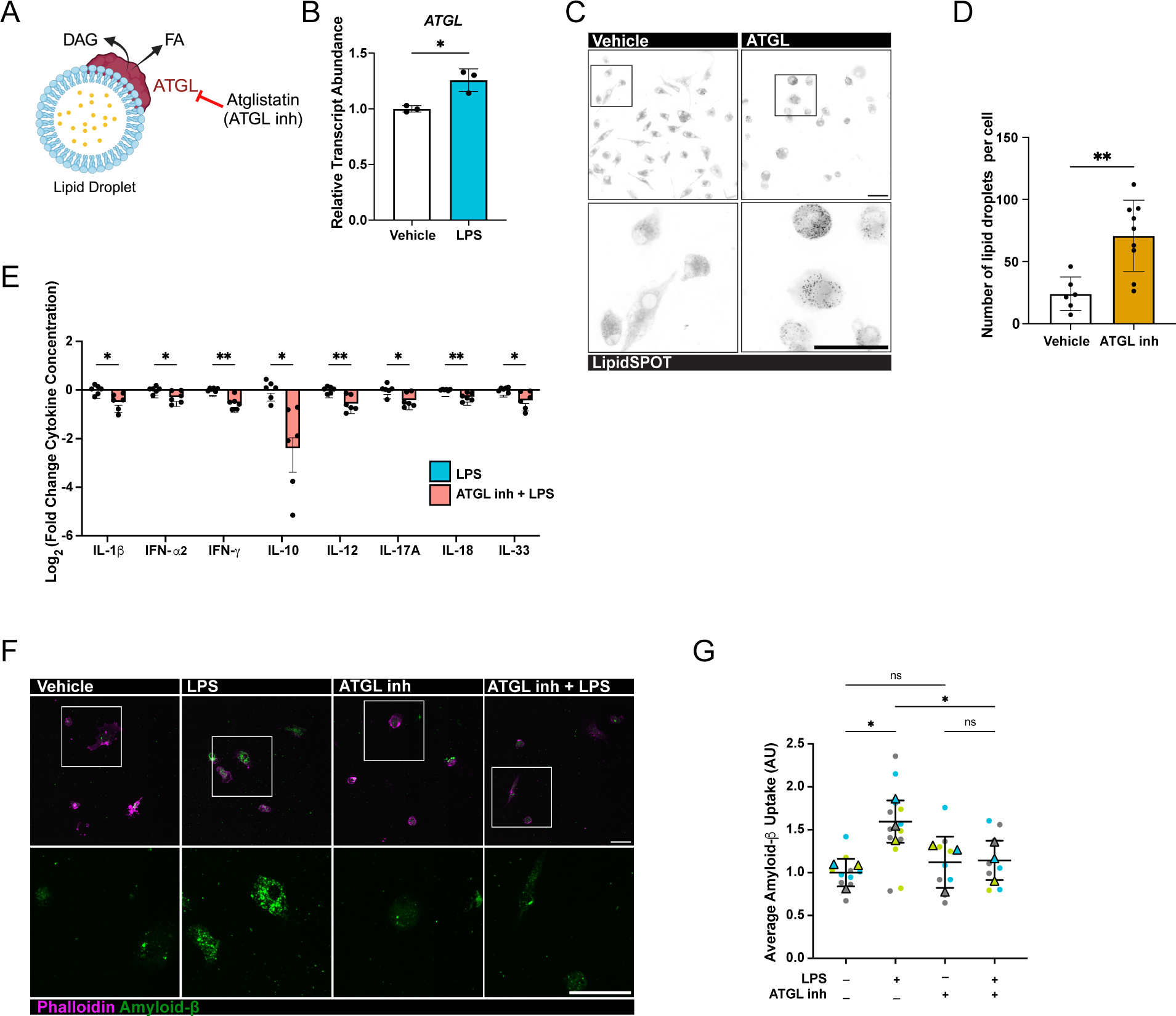
Utilization of LDs through catabolism necessary for LPS-mediated activation. **A.** Schematic depicting the inhibition of lipid droplet catabolism using atglistatin. **B.** Transcript abundance of adipose triglyceride lipase (*ATGL*) in LPS-treated microglia relative to vehicle-treated controls. (vehicle, white bars; LPS, blue bars; n = 3). * P ≤ 0.05 by unpaired t-test. **C.** Representative fluorescence images of *APOE3/APOE3* microglia treated with vehicle or ATGL inhibitor and stained with LipidSPOT. Bottom panels show high magnification of black box in the top panels. Scale bars: 50 μm. **D.** Quantification of the lipid droplet number per cell, with each dot representing an average of three wells with at least 20 cells analyzed. ** P ≤ 0.01 by unpaired t-test. **E.** Quantification of the fold change of cytokines secreted by lipopolysaccharide (LPS)-treated *APOE3/APOE3* microglia with (blue bars) and without inhibitor (red). n = 6 wells for 3 distinct biological cultures. * P ≤ 0.05; ** P ≤ 0.01; by unpaired t-tests. **F.** Representative fluorescence images of amyloid-beta (green) uptake assay in homeostatic and activated *APOE3/APOE3* microglia in the presence or absence of ATGL inhibitor. Cells were stained with Phalloidin (magenta). Bottom panels show high magnification of white box in the top panels. Scale bars: 50 μm. **G.** Quantification of amyloid-beta uptake (F) relative to vehicle-treated controls. Each dot represents an average of at least 5 cells per image. Each color represents an independent microglia differentiation. Triangles depict the average amyloid-beta uptake in each biological replicate. ns P > 0.05; * P ≤ 0.05; by one-way ANOVA, with post-hoc Šídák test to correct for multiple comparisons.

We then probed the effect of the ATGL inhibitor treatment on uptake of the disease-associated amyloid-beta peptide. We observed that, like DGAT inhibitor treatment, ATGL inhibitor treatment also inhibited the activation-associated uptake of amyloid-beta (Figure 4F,G), demonstrating a role for triglyceride catabolism in disease-associated processes. In combination with the DGAT inhibitor data presented earlier, these experiments establish that both the biosynthesis and catabolism of triglyceride-rich lipid droplets are necessary for microglia to fully respond to extrinsic stimuli.

### Modulating triglyceride biosynthesis controls immune state of *APOE4* microglia

One of the strongest risk factors for late-onset AD is *APOE4*. Our past work and other recent reports have all shown that *APOE4* microglia accumulate more lipid droplets than *APOE3* microglia^22,23^ even in the absence of any extrinsic stimuli. We confirmed that *APOE4* homozygous microglia contained more lipid droplets than their *APOE3* homozygous counterparts (Figure 5A,B). We also noticed that upon LPS activation, the lipid droplet level of microglia of both *APOE* genotypes were elevated to similar levels (Figure S5A,B).

**Figure 5.**
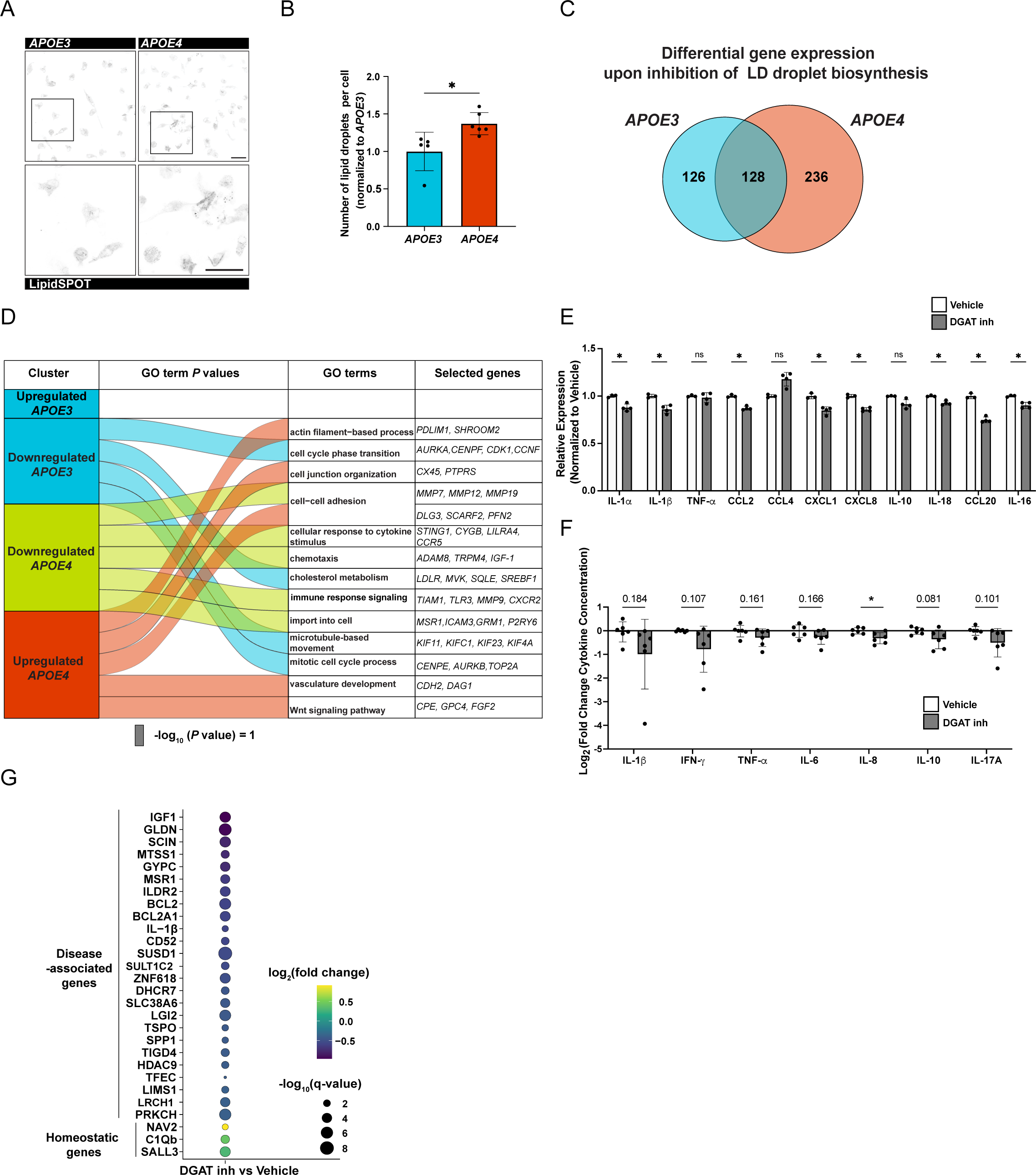
Modulating triglyceride flux controls the immune state of *APOE4* **microglia.** **A.** Representative fluorescence images of isogenic *APOE3/APOE3* and *APOE4/APOE4* human iPSC-derived microglia stained with LipidSPOT. Scale bar: 50 μm. Bottom panels show high magnification of images of regions of interest in top panels. **B.** Graph showing the quantification of the lipid droplet number per cell with each dot representing an average of at least 25 cells, in three wells analyzed. * P ≤ 0.05; by unpaired t-tests. **C.** Venn diagrams showing the overlap of the differentially expressed genes (DEGs) upon treatment with DGAT1 and DGAT2 inhibitors in *APOE3/APOE3* microglia and *APOE4/APOE4* microglia. **D.** Plot showing the overlap of the GO terms defined by DEGs from the *APOE3/APOE3* microglia and *APOE4/APOE4* microglia treated with DGAT1 and DGAT2 inhibitors relative to vehicle-treated controls. GO terms were determined using Metascape^57^. The width of the band for each GO term represents its P value calculated based on the cumulative hypergeometric distribution. The scale bar represents log_10_ (P value) = 1. Selected down- and up-regulated DEGs are listed for each GO term. **E.** Quantification of the expression levels of cytokines (from normalized read counts) relative to average vehicle-treated controls (vehicle, white bars; DGAT1 and DGAT2 inhibition, grey bars; n = 3). ns P > 0.05; * P ≤ 0.05 by unpaired t-tests. **F.** Quantification of the fold change of cytokines secreted by *APOE4/APOE4* microglia treated DGAT1 and DGAT2 inhibitors (grey bars) or vehicle (white bars) n = 6 wells across 3 distinct biological cultures. ns P > 0.05; * P ≤ 0.05; by unpaired t-tests. **G.** Dot plot showing the differential gene expression results for homeostatic and disease-associated microglia (DAM) marker genes identified across multiple studies. Color indicates log_2_ of the fold change of gene expression in DGAT1 and DGAT2-inhibitor treated *APOE4/APOE4* microglia compared to vehicle-treated. The size of the dot indicates the false discovery rate adjusted p-value.

After finding that modulation of triglyceride metabolism could alter LPS-mediated activation in *APOE3* microglia, we investigated the consequences of these modulators in *APOE4* microglia. To explore these effects in a controlled manner, we generated microglia from iPSC lines that were isogenic at all loci except *APOE* where they were either homozygous for *APOE3* or *APOE4*^21^ (Figure S1A). We treated both our *APOE3* and *APOE4* isogenic hiPSC-derived microglia with DGAT1 and DGAT2 inhibitors in the absence of any other activating stimuli and then profiled the transcriptomic pathways that were impacted by the inhibition of triglyceride biosynthesis. When we examined the transcriptome, we found that DGAT inhibitor treatment led to the differential regulation of ∼50% more genes in *APOE4* microglia than in their *APOE3* isogenic counterparts (Figure 5C). While DGAT inhibitor treatment predictably led to downregulation of mostly lipid biosynthesis pathways in *APOE3* microglia, in *APOE4* microglia, DGAT inhibitor treatment also led to a downregulation of immune pathways like cellular response to cytokine stimulus and immune response signaling (Figure 5D). Among the genes upregulated in *APOE4* microglia by DGAT inhibitor treatment, we found negative regulators of NF-κB -mediated inflammation like PDLIM1^36^, suggesting that DGAT inhibitor treatment modulates immune signaling in *APOE4* microglia. When specifically examining the expression of immune genes, we noticed that DGAT inhibitor treatment downregulated many cytokine and chemokine genes in *APOE4* microglia (Figure 5E). In line with our transcriptional findings, we also discovered that DGAT inhibitor treatment downregulated the secretion of inflammatory cytokines in *APOE4* microglia but not *APOE3* microglia (Figure 5F, S5C, S2H). These results demonstrate that even in the absence of any extrinsic stimulation, *APOE4* microglia share activation pathways similar to extrinsically activated *APOE3* microglia.

To determine whether gene expression changes were a result of chromatin accessibility changes, we performed ATAC sequencing on our *APOE4* microglia in the presence and absence of DGAT inhibitor treatment. We observed that the DGAT inhibitor treatment-associated transcriptional changes were not accompanied by changes in chromatin accessibility at the loci of differentially expressed genes (Figure S5D, Sy.x=6.1), suggesting that DGAT inhibitor treatment acts downstream of the chromatin structure.

Microglia in different AD models display several disease-associated transcriptional signatures^1,3–7^. Given *APOE4*’s key role as an AD risk factor, we explored whether modulation of triglyceride biosynthesis impacts disease-associated transcriptional responses characterized in several AD-associated microglial studies. We first examined the transcriptome of *APOE3* microglia treated with LPS and noticed that LPS-mediated activation upregulated several DAM microglia genes and downregulated several homeostatic microglial genes (Figure S5E). In *APOE3* microglia, DGAT inhibitor treatment renormalizes the transcriptome by upregulating homeostatic genes and downregulating disease-associated genes (Figure S5E). In the context of *APOE4* microglia, we observed that inhibition of triglyceride biosynthesis, even without any extrinsic stimuli, decreased disease-associated gene expression and upregulated homeostatic gene expression (Figure 5G). This suggests that modulation of triglyceride levels impacts the disease-associated states of *APOE4* microglia.

## Discussion

Within the past decade microglia have emerged as key mediators of neurological disease risk and progression. Many studies have characterized microglial states and their functional consequences^1–7,32^. In parallel, glial lipid accumulation has been rediscovered as key pathology in several neurodegenerative and neurological disorders^13,14,24,32,37–39^. Lipid droplets have also emerged as key organelles in the context of the immune response to pathogens, aging, and cancer by both modulation of cellular lipids and proteins ^32,40–45,46,47^. Recent studies have established that the *APOE4* AD risk polymorphism is associated with increased lipid accumulation in multiple glial cell types including astrocytes, microglia, and oligodendrocytes, and that this lipid accumulation has various cell non-autonomous consequences^18,19,22–24,28^.

Our work builds on this foundation by establishing that triglyceride accumulation is necessary for microglial response to activation. We used multiple iPSC lines from different genetic backgrounds to demonstrate that modulating triglyceride metabolism can modify how and whether a microglial cell responds to external or intrinsic stimuli. Triglyceride synthesis and, to a lesser extent, degradation, are necessary for microglial activation in the context of both *APOE3* cells as well as those harboring a risk factor for late-onset AD, *APOE4.* In the context of *APOE4*, inhibiting triglyceride synthesis restores the expression of homeostatic genes and suppresses the activation of disease-associated genes, indicating that *APOE4* cells have a pathological rewiring of lipid metabolism. This has been suggested in correlative data from other studies which show that *APOE4* microglia have both a higher lipid burden and a more “activated” phenotype than *APOE3* microglia^19,20,23,24^. Changes to triglyceride metabolism can also modulate essential microglial functions like phagocytosis and endocytosis of multiple substrates. Therefore, we propose that controlling the triglyceride content and storage in microglia is a promising avenue for controlling their inflammatory signaling and disease-associated functions.

In our study, we used stimulation of microglia with lipopolysaccharide as the primary paradigm to study microglial responses. LPS is traditionally used to mimic exposure to bacteria, however many of the genes and cytokines upregulated in response to LPS are similarly altered in AD (Figure S5E). Additionally, toll-like-receptor 4 (TLR4), which signals LPS-mediated activation, has been shown to be activated in a number of other inflammatory paradigms including neurodegeneration and high-fat diet^48,49^. By using LPS instead of amyloid-beta to trigger the inflammatory response, we were able to dissect the effects of lipid biology on the inflammatory response and on disease-associated amyloid uptake. Interestingly, another recent study identified DGAT inhibition to alleviate amyloid-beta induced phagocytosis deficits in mice^15^ suggesting that the same pathways we examined may be appropriate for modulating microglial response to not only LPS but other disease-associated stimuli as well.

We wondered the mechanism(s) by which controlling triglyceride metabolism impacts microglial response to immune stimuli. One possibility is that the triglyceride-rich lipid droplets store precursors for inflammatory lipids, a phenomenon that has been observed in multiple organisms^45,50,51^. In our microglia, we observe that the inhibition of triglyceride synthesis in the context of LPS activation results in a transcriptional downregulation or prostaglandin synthase (*PTGES*) (Figure S2C). Another possibility is that changes to triglyceride flux alter the lipid droplet proteome, which could include inflammatory factors some of which have been observed to be docked at lipid droplets in other cell types (like IGTP, IIGP1, and IFI47)^43,46^. Future studies will delve into the mechanisms of how triglyceride metabolism pathways control inflammation.

The inhibition of triglyceride biosynthesis and catabolism present a potential avenue for modulating disease-associated neuroinflammation for a variety of neurological conditions. Antisense oligonucleotides against DGAT1 are in clinical trials for fatty liver disease^52^. Inhibition of DGAT enzymes has also been suggested as therapeutic avenue for indications such as diabetes and glioblastomas^53,54^. However, there are many hurdles to overcome before DGAT inhibition becomes a viable therapeutic or preventative option. These include achieving delivery across the blood brain barrier (a challenge which has promising solutions in antibody-mediated delivery^55^) and correct dosing to avoid toxicity due to inhibition of the key fat storage functions of DGAT enzymes. This latter challenge could perhaps be overcome by exploiting functional redundancy of DGAT1 and DGAT2.

Our work has established a direct relationship between lipid storage and immune activity in microglia as well as shown that this process is hijacked by risk genotypes like *APOE4*. Future work can go into establishing the point in a disease course when immunomodulation has its greatest effect on progression. Our studies present new opportunities for tailoring microglial responses based on tuning their lipid state.

## Author Contributions

RAS: Conceptualization, methodology, investigation, validation, formal analysis, writing, visualization. KRJ: Methodology, software, formal analysis, editing. LC: methodology, validation. LGY: Methodology, investigation, project administration, JTR: software, data curation, visualization. JG: investigation, resources. HYS: investigation, formal analysis, supervision. PN: Conceptualization, methodology, validation, formal analysis, writing, visualization, supervision, funding acquisition.

**Supplemental Figure 1.**
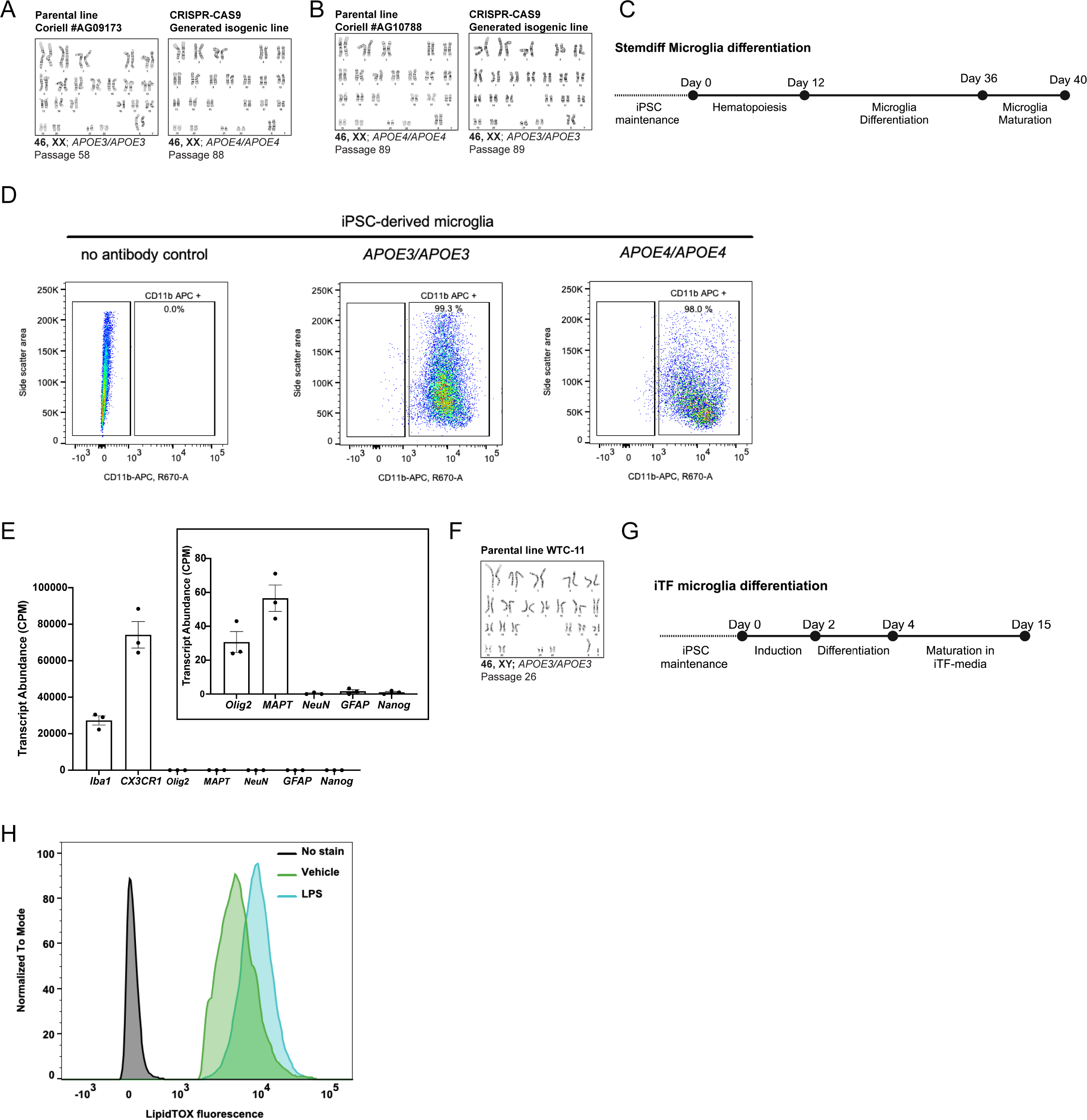
**A.** A normal karyotype was confirmed for parental line *APOE3/APOE3* Coriell #AG09173 and the isogenic *APOE4/APOE4* line. Karyotyping was done prior to microglia differentiation. **B.** A normal karyotype was confirmed for parental line *APOE4/APOE4* Coriell #AG10788 and the isogenic *APOE3/APOE3* line. Karyotyping was done prior to microglia differentiation. **C.** Overview of the microglia differentiation protocol to generate induced pluripotent stem cell (iPSC)-derived microglia using StemCell Technologies microglial differentiation workflow. **D.** Plots show the fluorescence-assisted cell cytometry analysis of *APOE3/APOE3* or *APOE4/APOE4* iPSC-derived microglia stained with anti-CD11b antibody or an unlabeled control. **E.** Expression (counts per million) of markers for microglia (Iba1, CX3CR1) and non-microglial genes. Zoom of graph panel shows the expression for markers of oligodendrocytes (Olig2), neurons (MAPT, NeuN), astrocytes (GFAP), and iPSCs (Nanog). **F.** A normal karyotype was confirmed for parental line WTC-11, induced-transcription (iTF) factor iPSCs prior to iTF-microglia differentiation. **G.** Strategy for generating iTF-Microglia from WTC-11 iTF-iPSCs. **H.** Quantification of lipid droplet levels (LipidTOX fluorescence) in WTC11 iTF-Microglia treated with LPS (blue) or vehicle control (green), analyzed by flow cytometry.

**Supplemental Figure 2.**
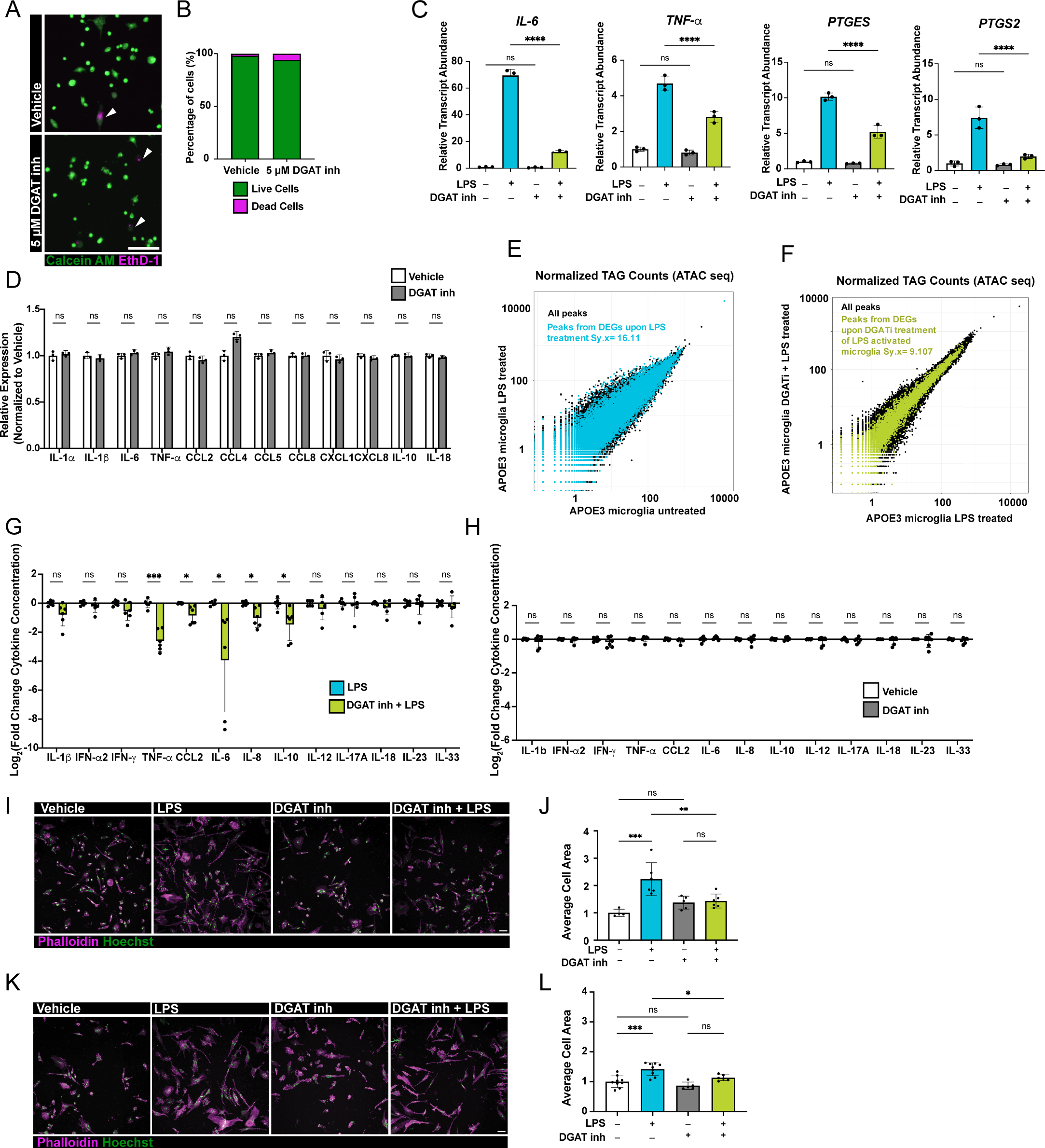
**A.** Merged representative fluorescence images of human iPSC-derived microglia stained with Calcein AM (green) and Ethidium homodimer-1 (EthD-1, magenta) to identify live and dead cells, respectively. Arrows indicate dead cells labelled by EthD-1. Scale bar: 100 μm. **B.** Quantification of the percentage of live (green bars) or dead (magenta bars) microglia (A) after treatment with vehicle or DGAT inhibitors. **C.** Transcript abundance of IL-6, TNF-α, Prostaglandin E synthase (PTGES), Prostaglandin-endoperoxide synthase 2 (PTGS2) in homeostatic and activated *APOE3/APOE3* microglia in the presence or absence of DGAT1 and DGAT2 inhibitors. (Vehicle, white; LPS-treated, blue; DGAT1 and DGAT2 inhibition, gray; DGAT1 and DGAT2 inhibition + LPS; green; n = 3). ns P > 0.05; **** P ≤ 0.0001 by one-way ANOVA, with post-hoc Šídák test to correct for multiple comparisons. **D.** Quantification of the expression levels of cytokines (from normalized read counts) relative to average vehicle-treated controls (vehicle, white bars; DGAT1 and DGAT2 inhibition, grey bars; n = 3). ns P > 0.05 by unpaired t-tests. **E.** Distribution of normalized tag counts (in bps) of all ATAC-seq detected peaks (black) and those corresponding to differentially expressed genes under LPS treatment (cyan). Sy.x was calculated using a linear regression model. **F.** Distribution of normalized TAG counts (in bps) of all ATAC-seq detected peaks (black) and those corresponding to differentially expressed genes under DGAT inhibitor treatment in the context of LPS activation (green). Sy.x was calculated using a linear regression model. **G.** Quantification of the fold change of cytokines secreted by lipopolysaccharide (LPS)-treated *APOE3/APOE3* microglia with (green) and without (blue bars) DGAT1 and DGAT2 inhibitors. n = 6 wells across 3 distinct biological cultures. ns P > 0.05; * P ≤ 0.05; *** P ≤ 0.001 by unpaired t-tests. **H.** Quantification of the fold change of cytokines secreted by *APOE3APOE3* microglia treated DGAT1 and DGAT2 inhibitors (grey bars) or vehicle (white bars). Fold change was calculated relative to vehicle controls. n = 6 wells across 3 distinct biological cultures. ns P > 0.05 by unpaired t-tests. **I.** Representative fluorescence images of homeostatic and activated microglia (*APOE3/APOE3* Coriell #AG09173) in the presence or absence of DGAT1 and DGAT2 inhibitors. Cells were stained with Phalloidin (magenta) and Hoechst 33258 (green). Scale bars: 50 μm. **J.** Quantification of the average cell area to assess microglia morphology (H). (vehicle, white; LPS-treated, blue; DGAT1 and DGAT2 inhibition, gray; DGAT1 and DGAT2 inhibition + LPS; green) Each dot represents an average of at least 385 cells across three wells analyzed. ns P > 0.05; ** P ≤ 0.01; *** P ≤ 0.001 by one-way ANOVA, with post-hoc Šídák test to correct for multiple comparisons. **K.** Representative fluorescence images of homeostatic and activated microglia (*APOE3/APOE3* edited from the parental line Coriell #AG10788) in the presence or absence of DGAT1 and DGAT2 inhibitors. Cells were stained with Phalloidin (magenta) and Hoechst 33258 (green). Scale bars: 50 μm. **L.** Quantification of the average cell area to assess microglia morphology (J). Vehicle, white; LPS-treated, blue; DGAT1 and DGAT2 inhibition, gray; DGAT1 and DGAT2 inhibition + LPS; green; Each dot represents an average of at least 377 cells across three wells analyzed. ns P > 0.05; * P ≤ 0.05; *** P ≤ 0.001 by one-way ANOVA, with post-hoc Šídák test to correct for multiple comparisons.

**Supplemental Figure 3.**
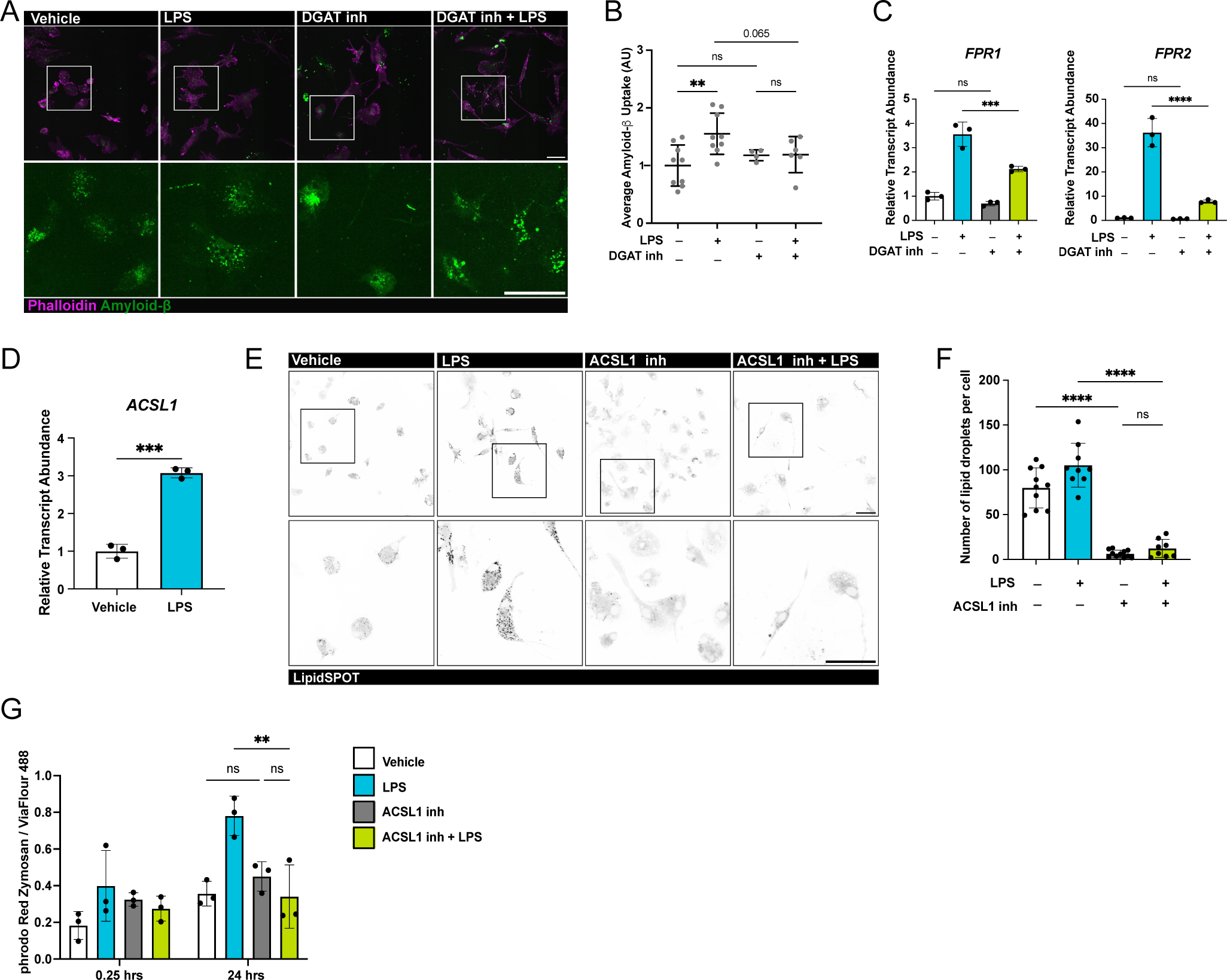
**A.** Representative fluorescence images of amyloid-beta (green) uptake assay in homeostatic and activated *APOE3/APOE3* microglia in the presence or absence of DGAT1 and DGAT2 inhibitors. Cells were stained with Phalloidin (magenta). Scale bars: 50 μm. Bottom panels show higher magnification of white box in top panels. (*APOE3/APOE3* edited from the parental line Coriell #AG10788) **B.** Quantification of amyloid-beta uptake (A) relative to vehicle-treated controls. Each dot represents an average of at least 20 cells per image. ns P > 0.05; ** P ≤ 0.01; by one-way ANOVA, with post-hoc Tukey test to correct for multiple comparisons. **C.** Transcript abundance of formyl peptide receptors (FPR1 and FPR2) in homeostatic and activated *APOE3/APOE3* microglia in the presence or absence of DGAT1 and DGAT2 inhibitors. (Vehicle, white; LPS-treated, blue; DGAT1 and DGAT2 inhibition, gray; DGAT1 and DGAT2 inhibition + LPS; green; n = 3). ns P > 0.05; *** P ≤ 0.001; **** P ≤ 0.0001 by one-way ANOVA, with post-hoc Šídák test to correct for multiple comparisons. **D.** Transcript abundance of Acyl CoA synthetase-1 (*ASCL1*) in vehicle- and LPS-treated microglia relative to vehicle-treated controls. (vehicle, white bars; LPS, blue bars; n = 3). *** P≤ 0.001 by unpaired t-test. **E.** Representative fluorescence images of homeostatic and activated *APOE3/APOE3* microglia stained with LipidSPOT, after treatment with ACSL inhibitor. Scale bars: 50 μm. Bottom panels show higher magnification of black outline in top panels. **F.** Quantification of the lipid droplet number per cell (E), with each dot representing an average of at least 10 cells, across three wells analyzed. (Vehicle, white; LPS-treated, blue; ACSL1 inhibition, gray; ACSL1 inhibition + LPS; green) ns P > 0.05; **** P ≤ 0.0001 by one-way ANOVA and post-hoc Šídák test to correct for multiple comparisons. **G.** Quantification of the fluorescence intensity of pHrodo Red Zymosan uptake normalized to ViaFlour 488 cytoplasmic stain per cell at time = 0.25 hours and time = 24 hours (Vehicle, white; LPS-treated, blue; ACSL1 inhibition, gray; ACSL1 inhibition + LPS; green; n = 3 wells). ns P > 0.05; ** P ≤ 0.01by one-way ANOVA, with post-hoc Šídák test to correct for multiple comparisons.

**Supplemental Figure 4.**
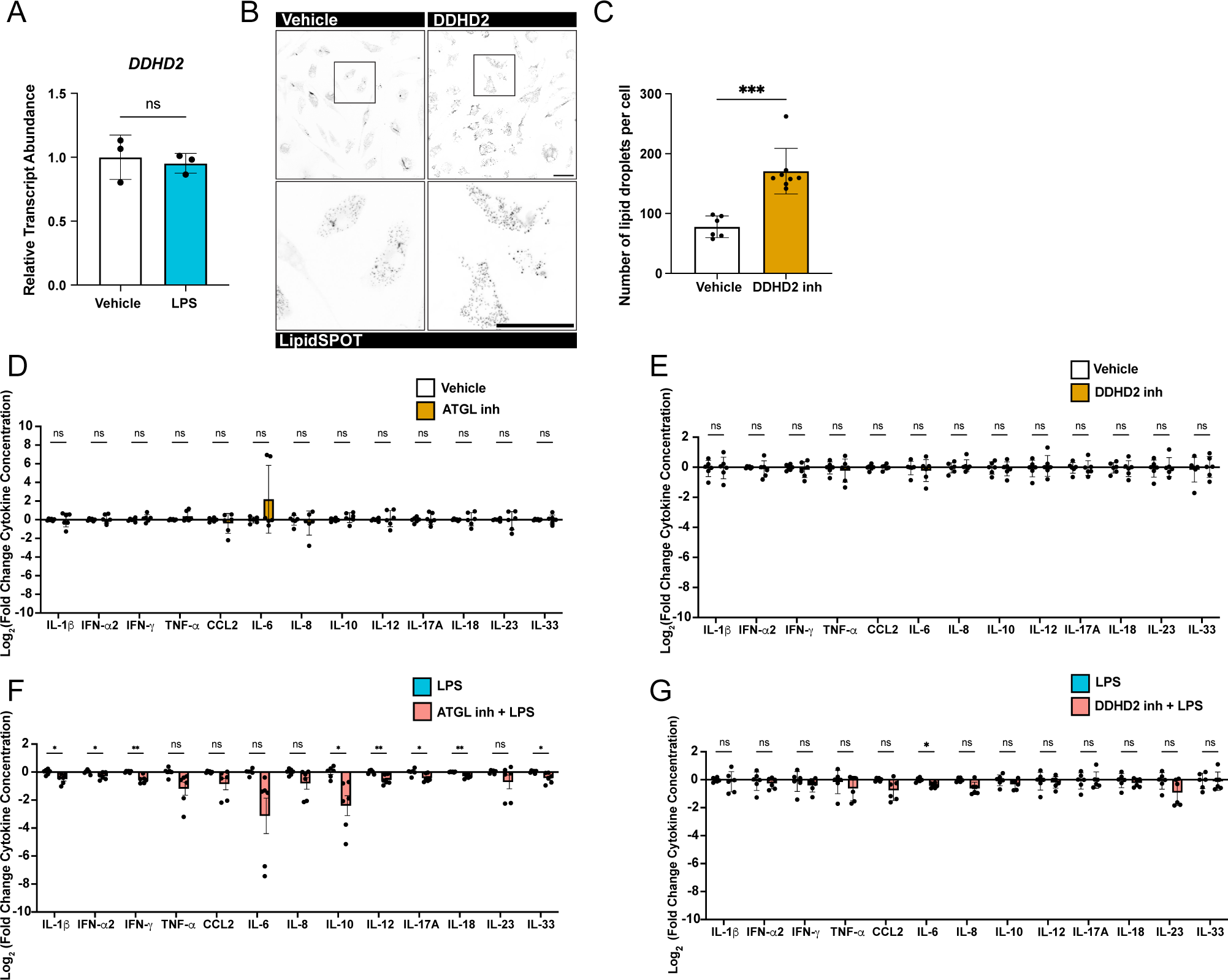
**A.** Transcript abundance of DDHD-domain-containing 2 (DDHD2) in vehicle- and LPS-treated microglia relative to vehicle-treated controls. (vehicle, white bars; LPS, blue bars; n = 3). ns P > 0.05 by unpaired t-test. **B.** Representative fluorescence images of homeostatic and activated *APOE3/APOE3* microglia stained with LipidSPOT, after treatment with DDHD2 inhibitor. Scale bars: 50 μm. Bottom panels show higher magnification of black outline in top panels. **C.** Quantification of the lipid droplet number per cell (B), with each dot representing an average of at least 11 cells, across three wells analyzed. (vehicle, white; DDHD2 inhibition, orange) *** P ≤ 0.001by unpaired t-test. **D.** Quantification of the fold change of cytokines secreted by *APOE3/APOE3* microglia treated ATGL inhibitor (orange bars) or vehicle (white bars). Fold change was calculated relative to vehicle controls. n = 6 wells across 3 distinct biological cultures. ns P > 0.05 by unpaired t-tests. **E.** Quantification of the fold change of cytokines secreted by *APOE3APOE3* microglia treated DDHD2 inhibitor (orange bars) or vehicle (white bars). Fold change was calculated relative to vehicle controls. n = 6 wells across 3 distinct biological cultures. ns P > 0.05 by unpaired t-tests. **F.** Quantification of the fold change of cytokines secreted by lipopolysaccharide (LPS)-treated *APOE3/APOE3* microglia with (red) and without (blue bars) ATGL inhibitor. Fold change was calculated relative to LPS-treated microglia. n = 6 wells across 3 distinct biological cultures. ns P > 0.05; * P ≤ 0.05; ** P ≤ 0.01 by unpaired t-tests. **G.** Quantification of the fold change of cytokines secreted by lipopolysaccharide (LPS)-treated *APOE3/APOE3* microglia with (red) and without (blue bars) DDHD2 inhibitor. Fold change was calculated relative to LPS-treated microglia. n = 6 wells across 3 distinct biological cultures. ns P > 0.05; * P ≤ 0.05; by unpaired t-tests.

**Supplemental Figure 5.**
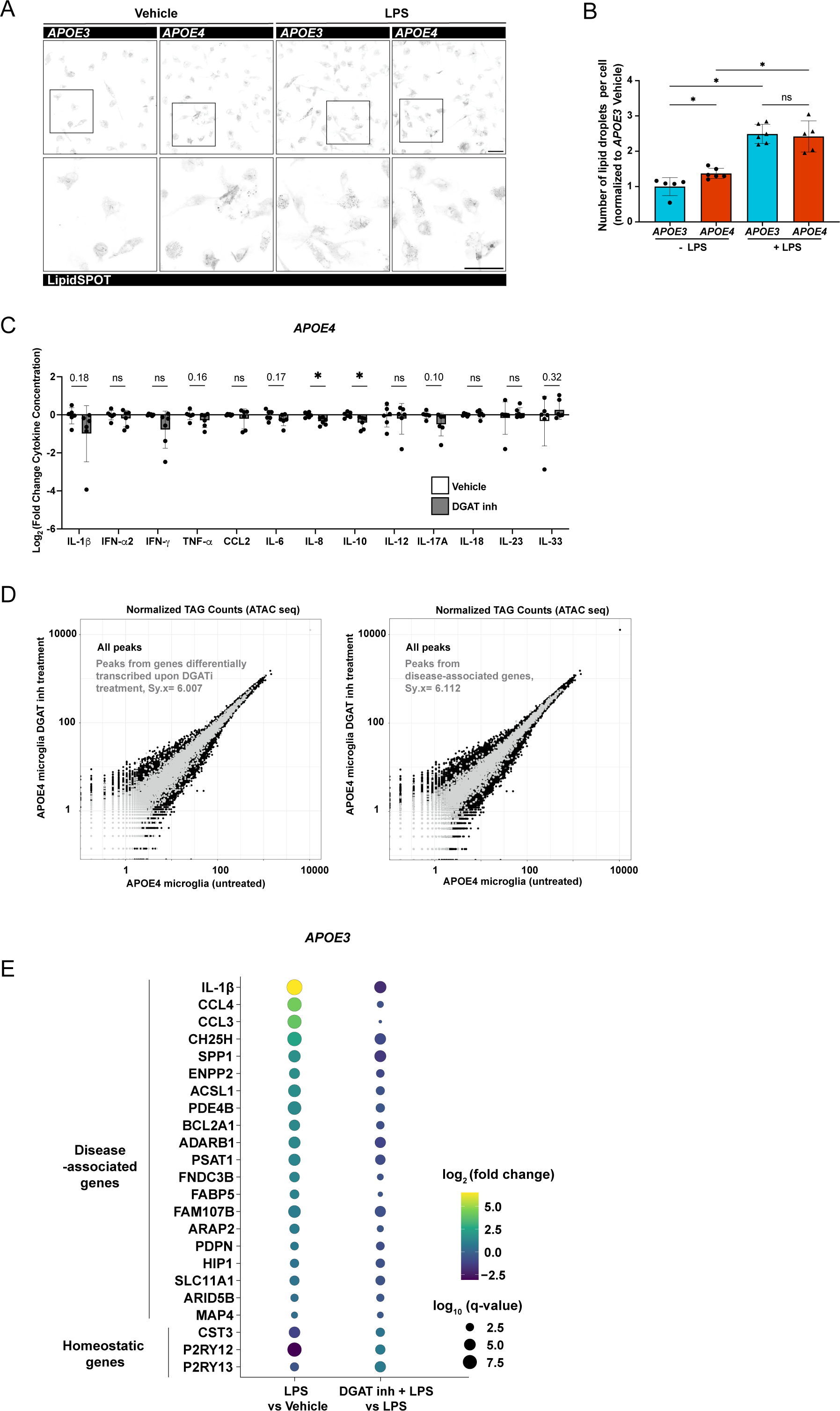
**A.** Representative fluorescence images of isogenic *APOE3/APOE3* and *APOE4/APOE4* human iPSC-derived microglia stained with LipidSPOT. Scale bar: 50 μm. Bottom panels show high magnification of images of regions of interest in top panels. **B.** Graph showing the quantification of the lipid droplet number per cell with each dot representing an average of at least 25 cells, in three wells analyzed. ns P > 0.05; * P ≤ 0.05; by one-way ANOVA, with post-hoc Šídák test to correct for multiple comparisons. **C.** Quantification of the fold change of cytokines secreted by *APOE4APOE4* microglia treated DGAT1 and DGAT2 inhibitors (grey bars) or vehicle (white bars). Fold change was calculated relative to vehicle controls. n = 6 wells across 3 distinct biological cultures. ns P > 0.05; * P ≤ 0.05 by unpaired t-tests. **D.** Distribution of normalized tag counts of all ATAC-seq detected peaks (black) and those corresponding to differentially expressed genes under DGAT inhibitor treatment (gray, left) or disease-associated genes (gray, right). Sy.x was calculated using a linear regression model. **E.** Dot plot showing the differential gene expression results for homeostatic and disease-associated microglia marker genes identified across multiple studies. Color indicates log_2_ of the fold change of gene expression of the indicated comparisons in *APOE3/APOE3* microglia. The size of the dot indicates the false discovery rate adjusted p-value.

## Materials and Methods

### iPSC maintenance

Two sets of isogenic *APOE3/APOE3* and *APOE4/APOE4* iPSCs (from parental lines Coriell #AG09173 and #AG10788, obtained via MTA from MIT) were thawed in a 6-well plate coated with hESC-qualified Matrigel Matrix (Corning) in Essential 8 Medium (Thermo) with 10 μM Y-27632 compound (StemCell Technologies). The media was replaced with Essential 8 Medium the following day and cultured until cells reached ∼90% confluence. Cells were passaged using *ReLeSR (*StemCell Technologies) for dissociation, as per manufacturer protocols.

### Karyotyping

iPSCs were cultured on hESC-qualified Matrigel Matrix-coated T25 cell culture flasks in Essential 8 Medium (ThermoFisher Scientific) until 60% confluence. They were then sent to WiCell Research Institute (Madison, WI) for G-banded karyotyping (20 metaphases equivalent) to ensure that each clone used for differentiations had no clonal abnormalities or chromosomal aberrations.

### Differentiation of Microglia from iPSCs

#### StemCell Technologies Stemdiff Microglia Kit Differentiations

iPSC-derived microglia were differentiated using the StemCell Technologies recommended microglial differentiation workflow (unless otherwise noted), which is based on the published protocol for the differentiation process^33^, hESC-qualified Matrigel-coated cultureware was used. Briefly, hematopoietic progenitor cells were generated using the STEMdiff Hematopoietic kit (StemCell Technologies). In lieu of half media changes, 1 mL media additions were performed throughout the hematopoietic differentiation protocol. Hematopoietic progenitors were harvested and passaged for microglia differentiation using STEMdiff Microglia Differentiation Kit (StemCell Technologies). Cells were maintained in this media for 24 days, according to manufacturer’s guidelines. For the last 4 days, microglia progenitors were cultured in the STEMdiff *Microglia* Maturation Kit (StemCell Technologies) medium to mature the microglia. Flow cytometry (CD11+ APC, Miltenyi Biotec) and immunocytochemistry (antibodies, see below) were used to assess purity.

#### iTF microglia differentiation

iTF-Microglia were generated based on previously published protocols^34^. Parental line WTC-11 (UCSF, MTA obtained) iPSCs expressing six inducible transcription factors (MAFB, CEBPα, IRF, PU1, CEBPβ, IRF5) were dissociated using StemPro Accutase (Gibco) and plated in Induction Medium (Essential 8 Medium,10 μM Chroman1, MedChemExpress, and 2 μg/mL Doxycycline hydrochloride, Sigma) at 400,000 cells per well on double-coated [hESC-qualified Matrigel Matrix and Poly-L-Ornithine (Sigma)] 10 cm tissue-culture treated plates. On day 2, the media was replaced with a differentiation media composed of Advanced DMEM/F12 (Thermo Fisher Scientific), 1X GlutaMAX (Thermo Fisher Scientific), 2 μg/mL Doxycycline hydrochloride, 100 ng/mL Human IL-34 (PeproTech) and 10 ng/mL Human GM-CSF (PeproTech). After 2 days, cells were washed with PBS and iTF-Microglia media was added (Advanced DMEM/F12, 1X GlutaMAX, 2 μg/mL Doxycycline hydrochloride, 100 ng/mL Human IL-34, 10 ng/mL Human GM-CSF, 50 ng/mL Human M-CSF (PeproTech), 50 μM Mevalonate (Sigma), 50 ng/mL Human TGFB1 (PeproTech), and 1X Antibiotic-Antimycotic (Gibco).

For dissociation, iTF-Microglia were washed with PBS before adding TrypLE Express (Gibco). After a 10 min incubation at 37 °C, cells were diluted 1:3 in Advanced DMEM/F12. iTF-Microglia were pelleted at 220 *g* for 5 min before resuspending in iTF-Microglia media.

Each biological replicate was performed using independent microglia differentiations.

### Immunocytochemistry

Microglia were fixed with 4% paraformaldehyde (Electron Microscopy Sciences) diluted in PBS for 20 minutes. After washing with 0.2% Triton X-100 in PBS three times for 5 minutes each, microglia were blocked with 1% normal donkey serum + 5% BSA blocking buffer for one hour at room temperature. Cells were incubated in primary antibodies (prepared in blocking buffer) overnight at 4°C (1:1000 anti-Iba1 (Novus Biologicals), 10 µg/mL anti-CX3CR1 (Abcam)). Microglia were washed with PBS three times, followed by a 1-hour incubation in secondary antibody (diluted in blocking buffer, 1:1000, Alexa Fluor 488 donkey anti-goat (Invitrogen), 1:1000 Alexa Fluor 488 donkey anti-rabbit (Invitrogen). Following three washes, images were acquired with a Nikon CSU-W1 Spinning Disk Microscope using Nikon Elements Software.

### LPS treatment

Microglia were treated with 5 µg/mL LPS, *E. Coli* O111:B4 (Millipore Sigma) in spent medium for 16 hours at 37 °C.

### Lipid droplet analysis

#### Modulating Lipid droplet burden

To assess the number of lipid droplets (LD) in microglia, cells were plated in an ibiTreated μ-Plate 96 well plate (Ibidi) in STEMdiff *Microglia* Maturation Medium or iTF-Microglia Medium. To decrease LD accumulation cells were treated 16 hours with a combination of inhibitors of 5 μM PF-04620110 (DGAT1, Sigma) and 5 μM PF-06424439 (DGAT2, Sigma) or 1 μM Triacsin C (ACSL, Tocris Bio-Techne). To inhibit LD lipase activity, cells were treated with either 1.3 nM KLH45 (DDHD2, Cayman Chemical) or 10 μM Atglistatin (ATGL, Sigma) for 6 hours.

#### Lipid droplet staining

Cells were carefully washed with PBS and stained with LipidSpot (Biotium) or LipidTOX (Invitrogen) as per manufacturer’s instructions. Images were acquired with a Nikon CSU-W1 Spinning Disk Microscope using Nikon Elements Software.

### Fluorescence-activated cell sorting (FACS)

To assess purity of Stemdiff microglia, cells were collected and resuspended in maturation media. After counting, Stemdiff microglia were stained with CD11+ APC (Miltenyi Biotec) at a 1:50 concentration in maturation media. Cell sorting was performed on BD FACSAria Fusion cytometer.

To assess lipid droplet accumulation in iTF microglia, cells were washed with PBS and stained with LipidTOX (Invitrogen), as per manufacturer’s instructions. Cells were then detached from the surface of the plate with TrypLE™(Gibco) following the manufacturer’s guidelines. After complete detachment, the cell suspension was centrifuged at 220x*g* for 5 minutes. Flow cytometry was performed on a BD LSR Fortessa Analyzer to measure the fluorescence intensity of LipidTOX.

Unstained controls were used in all FACS analysis. Gating and analysis were performed using FlowJo version 10.9.0 (FlowJo LLC).

### Microglia cell area quantification

Microglia were seeded at a density of 3500 cells per well in a 96-well imaging plate (ibidi). Following treatment with lipid modifiers, cells were fixed (as described above) and then washed with PBS. To identify the total number of cells, microglia nuclei were stained with 1:10,000 Hoechst 33258 (Sigma). Microglia were stained with Alexa Fluor 647 Phalloidin (Invitrogen). Using Nikon Elements GA3 Analysis suite, the total phalloidin positive area was masked and quantified in each image. That area was divided by the number of nuclei per image to get an average cell area to quantitatively characterize the microglia morphology.

### Viability Assay

To assess the viability of microglia, the Live/Dead Viability/Cytotoxicity Kit, for mammalian cells (Invitrogen) was used, as per manufacturer’s instructions. The percent of live or dead cells was determined for cells treated with vehicle or 5 μM PF-04620110 (DGAT1, Sigma) and 5 μM PF-06424439 (DGAT2, Sigma).

### Transcriptomic analysis

Cells were seeded at a density of 500,000 cells per well in a 12-well tissue-culture treated plate. Microglia were treated with LPS, DGAT inhibitors, or DMSO vehicle as described above. Cells were harvested after treatments in 300 μL of TRIzol (Invitrogen) and RNA was isolated using Direct-zol RNA isolation kits (Zymo Research). Illumina library preparation was performed to isolate polyadenylated transcripts and libraries were sequenced on an Illumina NovaSeq6000 sequencer.

Fastq files were quality inspected using the FastQC (https://www.bioinformatics.babraham.ac.uk) and MultiQC (https://multiqc.info) tools. For reference mapping (hg38) and gene count enumeration, the nf-core RNA-Seq pipeline (v3.10.1) was used (https://nf-co.re/rnaseq) in conjunction with select parameters (--clip_r1 10 --clip_r2 10). Counts produced were imported into R (https://cran.r-project.org/) then within-sample normalized using the “cpm” function. These normalized counts were then pedestalled by 2 and Log_2_ transformed. Genes not observed to not have a value >1 for any sample were filter removed with counts for surviving genes cross-sample normalized using the “cyclicloess” procedure available as part of the “normalizeBetweenArrays” function. Post cross-sample normalization, outliers were inspected for and confirmed absent by covariance-based Principal Component Analysis (PCA) scatterplot using the “princomp” function (cor=F) and by correlation-based heat map using the “cor” and “pheatmap” functions.

To remove noise-biased expression, locally weighted scatterplot smoothing was applied to the cross-sample normalized expression for all genes by sample class (Coefficient of Variation∼mean expression) using the “lowess” function. Fits produced per sample class were then over-plotted and inspected using the “plot” function to identify the common low-end cross-sample normalized expression value where the relationship between mean expression (i.e., “signal”) and Coefficient of Variation (i.e., “noise”) grossly deviated from linearity. Cross-sample normalized expression values were then floored to equal this value if less, while expression for genes not observed greater than this value for at least one sample were discarded as noise-biased.

For genes not discarded, expression differences across sample classes were tested for using the one-factor Analysis of Variance (ANOVA) test under Benjamini– Hochberg (BH) False Discovery Rate (FDR) Multiple Comparison Correction (MCC) condition. For ANOVA, the “Anova” function was used. For correction, the “mt.rawp2adjp” function was used. Genes having a corrected P < 0.05 by this test were then subset and the Tukey Honest Significance Difference test used to generate mean differences and p-values for each possible pairwise comparison of sample classes. The function used to do so was “TukeyHSD”. Genes having a post-hoc P < 0.05 for a specific comparison and a linear difference of means >= 1.5X for the same comparison were deemed to have expression significantly different between the sample classes compared. Post testing, the union set of differential genes identified across sample comparisons were used to investigate sample-to-sample relationships via the same visuals and functions used to confirm the absence of outliers.

### Detection of NF-κB p65 nuclear localization

For fluorescence detection of NF-κB p65 nuclear localization, microglia were fixed, and the standard immunocytochemistry technique described above was carried out using 1:1000 anti-NFk-B (Cell Signaling Technology) and 1:1000 Alexa Fluor 568 donkey anti-rabbit (Invitrogen). Microglia nuclei were stained with Hoechst 33258. Fluorescence signal was captured using Nikon CSU-W1 Spinning Disk Microscope. Using Nikon Elements GA3 image processing suite, nuclei were segmented and the intensity of nuclear NF-κB p65 within each nuclear area was measured for each group.

### Detection and Quantification of Cytokines and Chemokines

Cells were seeded at a density of 500,000 cells per well in a 12-well tissue-culture treated plate. Microglia were treated with LPS, DGAT inhibitors, ATGL inhibitor, DDHD2 inhibitor or DMSO control (as described above).

#### Chemokine bead array

The Chemokine concentrations in cell culture media were assessed using the 13-plex LEGENDplex Human Inflammation Panel 1 (Biolegend), according to the manufacturer’s instructions. Samples were then run on a LSRFortessa Cell Analyzer (BD Biosciences) until at least 5,000 events had been recorded. Data were analyzed using Qognit Software (Biolegend). The bioactive IL-12p70 was abbreviated as IL-12 in figures.

### Uptake assays

Microglia were seeded at a density of 3500 cells per well onto fibronectin coated (StemCell Technologies) 96-well plates (PerkinElmer or Ibidi). To measure the phagocytic ability of microglia, three ligands were profiled: 70,000 MW Dextran (Invitrogen), amyloid-beta-42 labeled with HiLyte Fluor-555 peptide (AnaSpec) and pHrodo Zymosan BioParticles (Invitrogen).

#### Disease-relevant substrate uptake

To assess amyloid-beta-42 uptake by microglia-like cells, a 1 mg/mL stock solution of the peptide was prepared following the manufacturer’s guidelines. Microglia were then treated with 1 μg/mL amyloid-beta-42-555 for one hour at 37°C. Cells were then fixed using 4% paraformaldehyde and washed with PBS. For imaging, microglia were stained with Hoechst 33258 and Alexa Fluor 647 Phalloidin. 10 μm image z-stacks were captured and analyzed using Nikon Elements GA3 image processing suite, to measure the fluorescence intensities of amyloid-beta-42-555 within a cell.

#### Endocytosis

A 25 mg/mL stock solution of 70,000 MW Texas Red-Dextran was prepared according to the manufacturer’s guidelines. The uptake assay described above was repeated with 20 μg/mL Texas-Red Dextran.

#### Phagocytosis of pHrodo Zymosan BioParticles

Microglia were stained with 1 μM ViaFlour 488 SE (Biotium) in prewarmed PBS for 10 mins at 37°C. After staining with the cytoplasmic dye, the solution was replaced with 1X Live Cell Imaging Solution (Invitrogen). pHrodo Zymosan BioParticles were reconstituted according to manufacturer’s protocols. 20 μg/mL pHrodo Zymosan was prepared in 1X Live Cell Imaging Solution. Cells were then treated with the pHrodo zymosan solution and imaged over a 24-hour period using Satorius Incucyte Live-Cell Imaging System. This time series experiment consisted of four frames per well, with a time interval of one hour between each capture. The fluorescence intensities were analyzed using the Sartorius Analysis System. The amount of Zymosan BioParticles taken up by microglia was calculated using the area of red puncta within cells/the area of cytoplasmic dye.

### ATAC Sequencing

ATAC-seq was performed as described previously^58^. Briefly, 35,000-76,000 microglia were washed with 1xPBS, followed by accessible chromatin tagging and fragmentation with 50ul transposase cocktail (25 μl of 2x TD buffer, 2.5 μl of TDE1, 0.5 μl of 1% digitonin, 22 μl of nuclease-free water) (Illumina, #20034197 and Promega, #G9441) at 37°C for 30 minutes with 300 rpm shaking. The fragmented DNAs were purified using a QIAGEN MinElute kit (Qiagen, #28006). The ATAC-seq libraries were constructed and amplified using NEBNext High-Fidelity 2x PCR Master Mix (NEB, #M0541) and SYBR Green I (Thermo Fisher Scientific, #S-7563), followed by a QIAGEN PCR cleanup kit (Qiagen, #28106). The libraries were then sequenced on NovaSeq (Illumina) at the NIAMS sequencing core facility. The 50bp paired-end reads were mapped to the human genome hg19 using Bowtie, followed by removal of redundant reads using FastUniq. BigWig tracks were further created by HOMER and analyzed by IGV genome browser.

### Quantification and statistical analysis

Details of experiments can be found in figure legends. To determine statistical significance, GraphPad Prism 9 software was used to perform unpaired t-tests, one-way ANOVA, or two-way ANOVA, with post hoc analysis as indicated in each figure legend. Data are represented as mean ± SD; ns P > 0.05; * P ≤ 0.05; ** P ≤ 0.01; *** P ≤ 0.001; **** P ≤ 0.0001. FlowJo (version 10.9.0) was used to process fluorescence-activated cell sorting data. Imaging processing was performed using Nikon Elements GA3 image processing and Sartorius Analysis System.

All schematics were generated using BioRender.com. Final Venn diagram were generated using the Eulerr.co website (https://eulerr.co/). Hypergeometric tests were performed on the Graeber Lab website (https://systems.crump.ucla.edu/hypergeometric/). GO enrichment analyses were performed using ShinyGO 0.80 and Metascape. Graphs were made using GraphPad Prism v10. Alluvial plots and scatter plots, were generated with ggplot2 on R studio.

## Acknowledgements

We thank Artur Gevorgyan, Dr. Carla Cuni-Lopez, Linda Yang, and Dr. Richard Proia for critical reading of this manuscript. We would like to acknowledge Dr. Erika Lara and Dr. Victor Bass for their help in experimental setups. This work was funded by the Intramural Research Programs of the National Institute for Diabetes and Digestive and Kidney Diseases (RAS, LC, LY JTR, PN), the National Institute for Neurological Disorders and Stroke (RAS, KJ, LC, LY, JTR, JG, HYS, PN), and the National Eye Institute (JG, HYS). This work was also supported by the Center for Alzheimer’s Disease and Related Dementias (CARD) at the National Institute of Aging (PN) and the Chan Zuckerberg Initiative (PN). We sincerely thank the NHLBI Flow Cytometry Core (Dr. Pradeep Dagur, Dr. Maria Lopez-Ocasio, Dr. Steve Hockman, and Dr. Wan-Chi Lin), NIH Intramural Sequencing Core, and NHLBI Genomics core (Dr. Yuesheng Li, Dr. Yan Luo, and Dr. Patrick Burr) for their help with flow cytometry and sequencing.

The WTC-11 iPS cells were obtained through Dr. Martin Kampmann under MTA from UCSF. The AG09173 and AG10788 iPSC lines and derivatives were obtained from Dr. Li-Huei Tsai under MTA from MIT.

